# *Pseudomonas aeruginosa* co-opts host antibiotic metabolism by pyocyanin induction of cytochrome P450 enzymes

**DOI:** 10.64898/2026.07.25.740721

**Authors:** Emily G. Gracey, Sylvie E. Kandel, Jed N. Lampe

## Abstract

The human opportunistic pathogen *Pseudomonas aeruginosa* produces copious quantities of the secondary metabolite pyocyanin (PYO). PYO has recently been shown to interact with the aryl hydrocarbon receptor (AhR), a transcription factor controlling expression of a number of genes, including the cytochrome P450 CYP1A family involved in fluoroquinolone antibiotic clearance. In this study, we investigated whether *Pseudomonas aeruginosa* could influence the metabolism of ciprofloxacin through human *CYP1A2* induction. Primary human hepatocytes were exposed to 1-100 μM PYO, with high concentrations resulting in apparent cytotoxicity. Treatment with 5 µM PYO led to a 6.2-fold change in *CYP1A2* mRNA and a 3.6-fold increase in oxociprofloxacin metabolite formation. Our results suggest that sub-toxic concentrations of PYO induce *CYP1A2* expression and increase ciprofloxacin metabolism, which could lead to sub-efficacious antibiotic concentrations and further drive resistance. This finding directs us to a novel mechanism by which *Pseudomonas aeruginosa* may escape antimicrobial therapy by hijacking host xenobiotic metabolism pathways.

**GRAPHICAL ABSTRACT:** 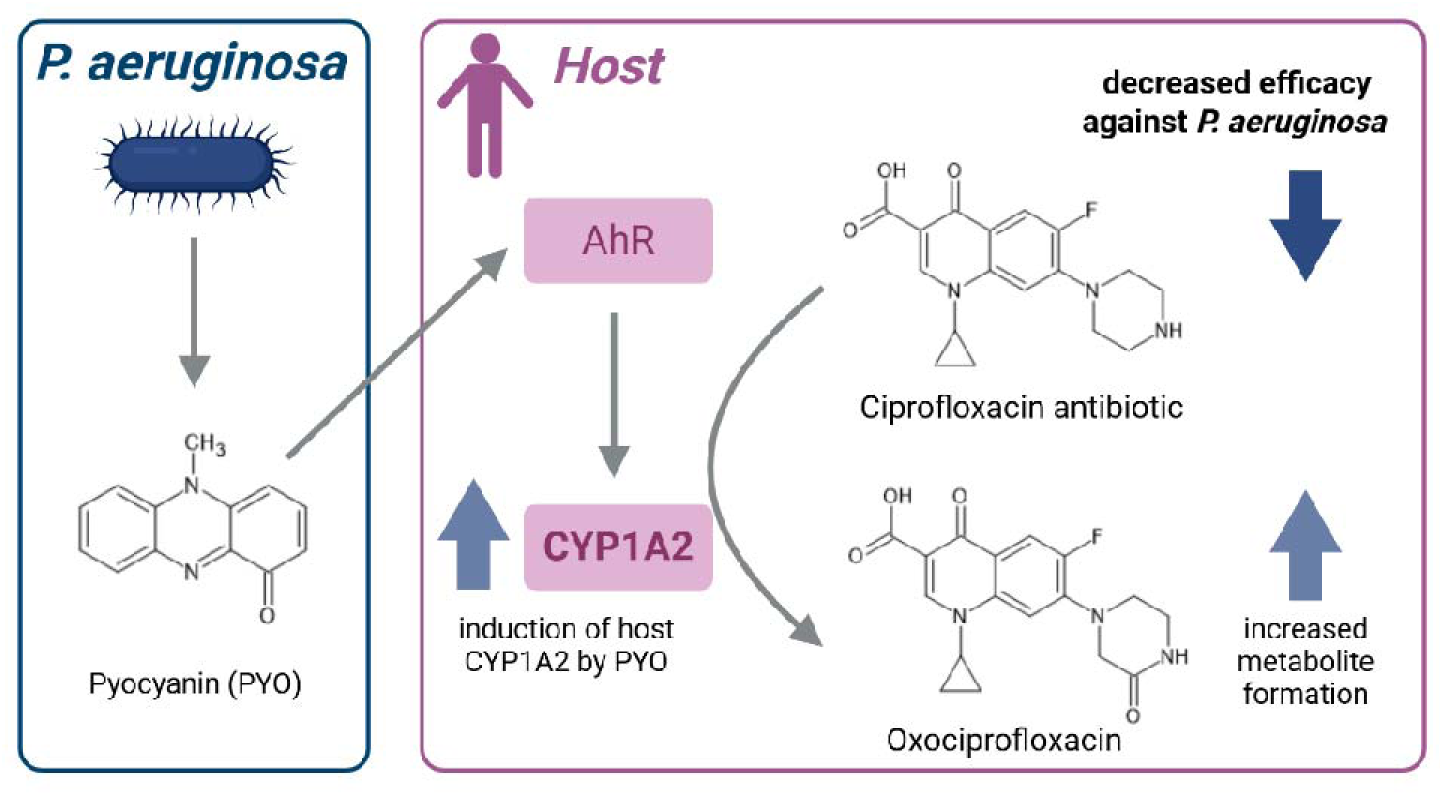

## INTRODUCTION

Pathogenic microorganisms have evolved a remarkable number of chemical strategies to induce the host organism to produce a hospitable environment in which to grow and replicate.^1, 2^ One mechanism by which they do this is by producing secondary metabolites that act as host cell toxicants facilitating entry of intercellular pathogen molecules into the host cell or also that may act by down regulating the host immune response.^3–5^ The human opportunistic pathogen *Pseudomonas aeruginosa* (*P. aeruginosa*) produces substantial quantities of the secondary metabolite pyocyanin (PYO), a blue phenazine (5-methylphenazin-1(5*H*)-one) dye contributing to *P. aeruginosa*’s characteristic blue-green color. PYO is a zwitterion at physiological pH and can readily cross both bacterial and eukaryotic cell membranes, serving a dual role as virulence factor and electron carrier for *P. aeruginosa*.^6^ PYO is cytotoxic and can reach concentrations of over 100 μM in sputum from infected patients,^7^ making the bioburden of this compound significant. In addition to its cytotoxicity, PYO has recently been shown to bind to the aryl hydrocarbon receptor (AhR),^8^ a ligand-dependent transcription factor and environmental probe that can activate the transcription of xenobiotic metabolizing enzymes, cytokines and chemokines, as well as regulate inflammatory leukocyte recruitment.^9, 10^ Modulation of the AhR may allow *P. aeruginosa* to alter the host immune response, as has previously been demonstrated with several of its other secondary metabolites,^11^ thereby providing a more permissive environment for pathogenic growth and proliferation. AhR is also important for the regulation of the expression of some drug metabolizing cytochrome P450 (CYP P450) enzymes,^12^ including CYP1A2 which is found constitutively expressed at high level in the human liver and inducible in the brain, pancreas, gastrointestinal tract, liver, and lung.^13^

One of the first line of antibiotics used to treat *P. aeruginosa* infection is the fluoroquinolone ciprofloxacin.^14, 15^ Despite its effectiveness *in vitro*, at least 20% of cystic fibrosis patients taking ciprofloxacin for *P. aeruginosa* lung infections remain unresponsive due to poor pharmacokinetics.^16^ While the exact source of this therapeutic recalcitrance is unknown, previous studies have indicated that it is *not* due to pharmacogenetic variation within the host.^16^ Interestingly, ciprofloxacin,^17^ is thought to undergo hepatic CYP1A2-mediated clearance, with two Phase 1 metabolites identified in human serum and urine samples: the desethyl- and oxo-ciprofloxacin derivatives.^18, 19^ Furthermore, previous studies have identified ciprofloxacin as a known inhibitor of CYP1A2 *in vivo.*^17^ Therefore, this raises the intriguing possibility that, through the production and secretion of PYO, *P. aeruginosa* may induce the expression of a host xenobiotic metabolizing enzyme that is responsible for clearing the very antibiotic that targets it. To test this hypothesis, primary human hepatocytes were treated with PYO, subsequently dosed with ciprofloxacin, and variations in *CYP1A2* expression level and ciprofloxacin metabolism were measured. Our results indicate that PYO is not only a potent inducer of hepatic *CYP1A2* expression, but that it also significantly increases the rate in ciprofloxacin metabolic oxidation.

## RESULTS & DISCUSSION

Pooled primary human hepatocytes were treated with PYO, and cellular response was measured in terms of gene expression, ciprofloxacin metabolism, and cytotoxicity as depicted in **Figure 1**. Upon treatment with PYO, a concentration-dependent increase of *CYP1A2* mRNA level was observed up to 5 μM (**Figure 2**). Fold change in *CYP1A2* mRNA compared to the dimethyl sulfoxide (DMSO) control (1.06 ± 0.42) was 2.40 ± 0.80 at 1 μM PYO, 6.17 ± 1.97 at 5 μM PYO, and 5.40 ± 1.37 at 20 μM PYO (**Figure 2A**). This semi-concentration dependent increase in *CYP1A2* expression was significant when compared with the DMSO control and mirrors the positive control, methylene blue (MB) (fold change 12.04 ± 2.44), a known CYP1A2 inducer, with a tricyclic core structure resembling PYO. This demonstrates that PYO can induce known drug metabolizing enzymes of the host in a physiologically relevant and concentration-dependent manner, likely acting through modulation of the AhR receptor.^12^ AhR has previously been shown to be sensitive to activation by bacterial metabolites, including polyphenolics and indoles.^8, 11^ It recognizes xenobiotic response elements that result in increasing the expression of *CYP1A1*, *CYP1A2*, and *CYP1B1*.^12^ However, CYP1A1 and CYP1B1 are thought to primarily play a role in endogenous functions, not xenobiotic detoxification.^20–22^ As the name suggests, the main ligands for AhR are planar polycyclic aromatic hydrocarbons, which also happen to be the preferred CYP1A2 substrates.^23^ The structure of PYO is consistent with this trend and there is some structural overlap with the ciprofloxacin substrate itself. While circulating plasma concentrations of PYO have not yet been reported, the lower concentrations used in our study represent 1/100^th^ and 1/20^th^ of the amount reported to be present in human sputum samples infected with *P. aeruginosa*,^7^ indicating that *CYP1A2* induction is observed even at relatively low PYO concentrations. *CYP3A4* expression was also tested as negative control, with significant increases only observed with rifampicin (RIF), a known CYP3A4 inducer operating via other transcription factors (supplemental **Figure S1**).^12^

**Figure 1.**
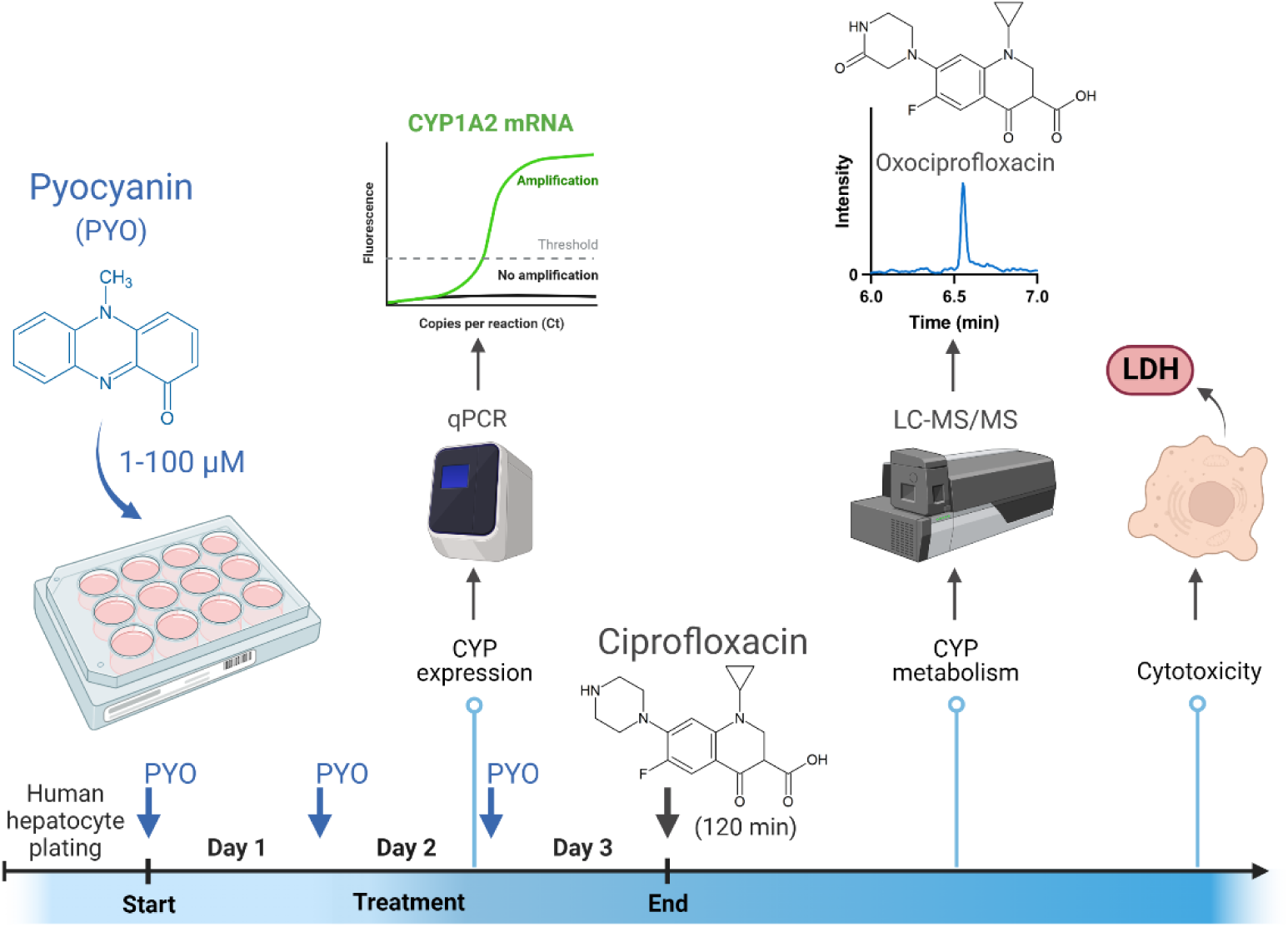
Experimental model schematic for assessing PYO effect on *CYP1A2* expression, ciprofloxacin metabolism, and cytotoxicity in primary human hepatocytes.

**Figure 2.**
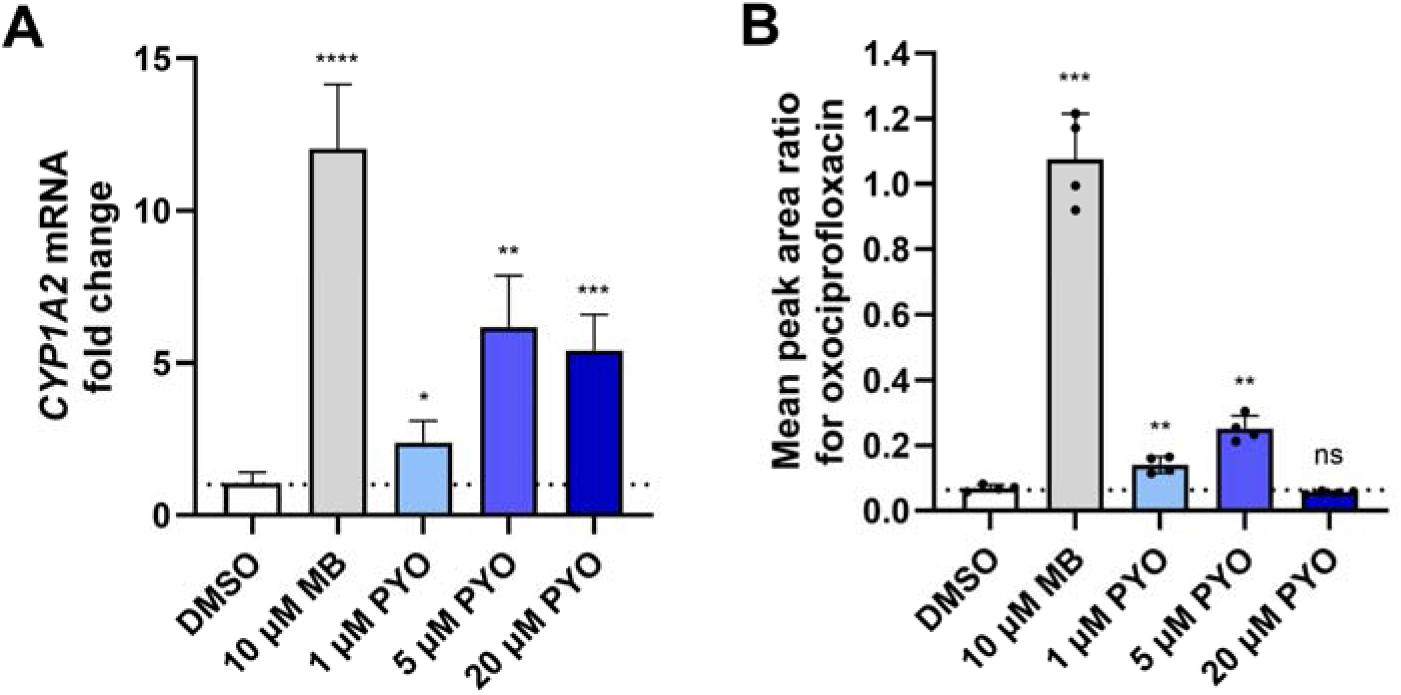
PYO effect on *CYP1A2* expression and oxociprofloxacin metabolite formation in plated pooled cryopreserved human hepatocytes. *A*, Fold change of *CYP1A2* mRNA levels in hepatocytes treated once daily for two days with medium containing either DMSO (solvent control), 10 µM methylene blue (MB; positive control) or PYO at 1, 5 or 20 µM. All four biological replicates were assayed by duplicate qPCR reactions. *B*, Fold change of peak area ratios for the oxociprofloxacin metabolite identified by LC-MS in the 120 min ciprofloxacin (30 µM) incubations done with plated human hepatocytes previously treated for up to three days with either DMSO, 10 µM MB or PYO at 1, 5 or 20 µM. The bar graphs represent the means of four biological replicates (individual data points shown) with error bars as standard deviations, for both the mRNA and activity data.

Hepatocyte treatment with PYO also led to a concentration-dependent increase in the metabolism of ciprofloxacin to oxociprofloxacin (**Figure 2B**). Oxociprofloxacin peak area ratio fold changes were 1.00 ± 0.14 for the DMSO control, 2.02 ± 0.36 at 1 μM PYO and 3.59 ± 0.56 at 5 μM PYO. A decrease in metabolite production was observed at 20 μM PYO (0.80 ± 0.07), possibly due to the emergence of cytotoxicity at this concentration. The oxociprofloxacin metabolite was positively identified using a known synthetic standard (**Figure 3**), and was the only metabolite found to increase in the PYO treated hepatocyte cultures, further implicating the involvement of CYP1A2 in the metabolism of ciprofloxacin. In humans, ciprofloxacin is metabolized into oxociprofloxacin, one of the primary urinary metabolites.^18, 24^ While fluoroquinolone antibiotics are typically effective, as a class they suffer from poor ADME (Absorption, Distribution, Metabolism, and Excretion) properties in part due to their chelating effects and variable pharmacokinetics, as noted above.^16, 25^ Hence, induction of the CYP1A2 enzyme could lead to increased clearance and reduced efficacy of the drug, ultimately promoting antibiotic resistance in the pathogen. Indeed, large pharmacokinetic differences in the disposition of ciprofloxacin exist across patient populations,^26^ including individuals with cystic fibrosis,^16, 27^ altogether indicating the presence of unidentified factors influencing the drug’s disposition. These differences can promote treatment failure and development of antibiotic resistance. This has implications beyond *P. aeruginosa* infections in the lung, as *P. aeruginosa* is also known to be a primary cause of nosocomial infections in burn victims^28^ and was recently reported to be the causative agent in eye infections due to contaminated saline drops that led to hospitalization and, in four cases, death.^29^ Furthermore, *P. aeruginosa*, as well as other infectious pathogens, are known to secrete a variety of secondary metabolites, from the phenazine class and others, that have not been examined for their ability to induce drug metabolizing P450 expression in relevant tissue systems.

**Figure 3.**
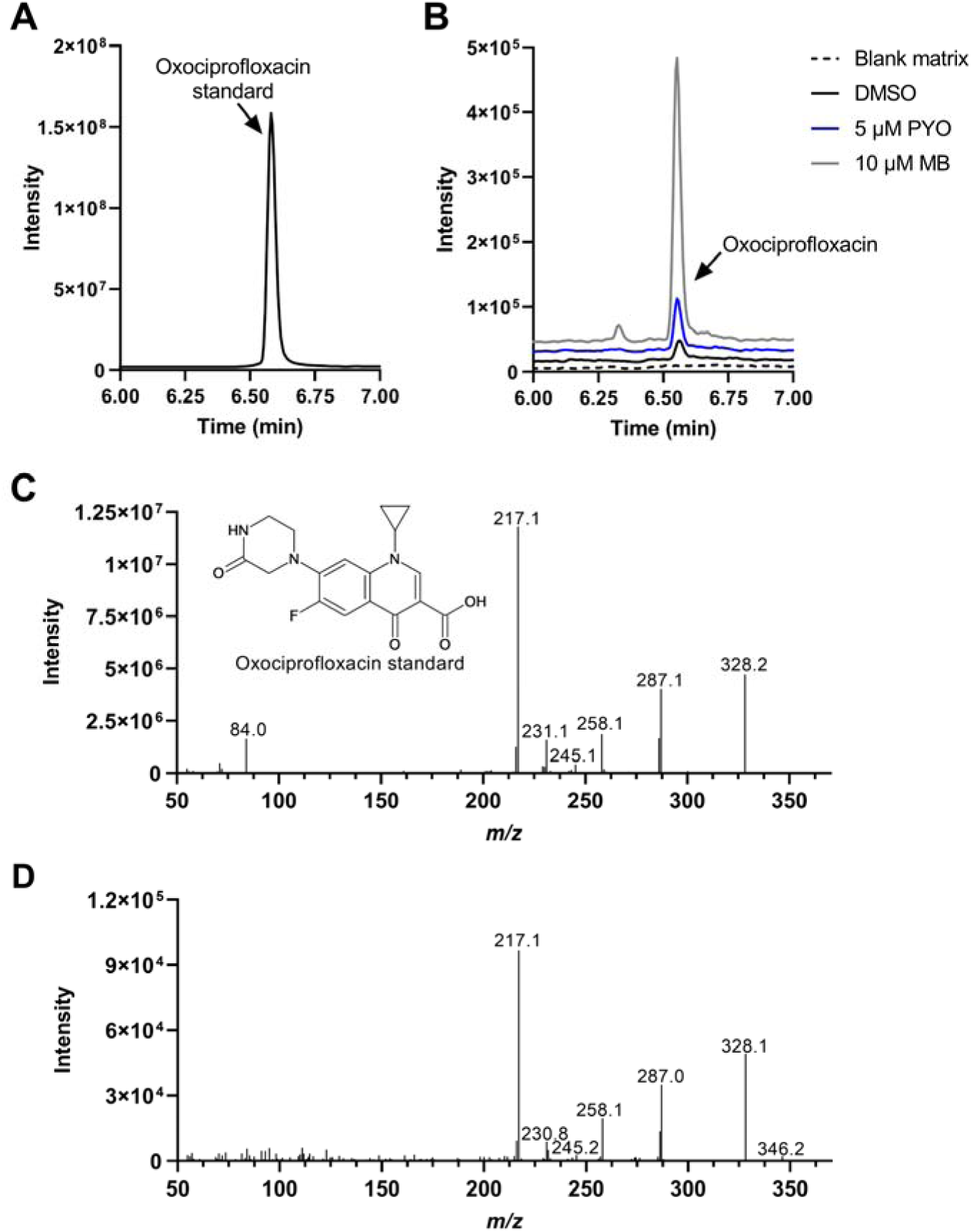
LC-MS analysis for the oxociprofloxacin metabolite produced in treated hepatocyte cells. *A*, Representative MRM chromatogram for the oxociprofloxacin standard at 0.5 µM in matrix. *B*, Representative MRM chromatograms for oxociprofloxacin in blank matrix and hepatocyte incubations after DMSO, 5 µM PYO or 10 µM MB treatment. *C*, MS/MS spectrum of the oxociprofloxacin standard. *D*, MS/MS spectrum of the oxidative metabolite detected in hepatocyte incubations with ciprofloxacin. Retention time and MS/MS spectrum matched the oxociprofloxacin standard.

Unsurprisingly, hepatocytes exposed to high concentrations of pyocyanin resulted in major cellular toxicity. The highest concentration of pyocyanin tested (100 μM) was highly toxic to the hepatocytes with a lactate dehydrogenase (LDH) release fold change of 7.05 ± 0.76 and led to cell death and detachment from the plate (**Figure 4**). A midrange concentration of 25 µM PYO showed an increase in LDH release (fold change of 3.04 ± 0.84) compared to the DMSO control. The cells were still attached to the plate but with a different morphology compared to the DMSO control. No significant changes in LDH release were observed with either controls (DMSO or MB) or PYO treatment between 1 and 20 μM (supplemental **Figure S2**); therefore this is the PYO concentration range assessed for *CYP1A2* mRNA and activity. The onset of cytotoxicity ∼ 20-25 μM PYO could explain the decrease in ciprofloxacin metabolism, where the cells were unhealthy and becoming metabolically incompetent, but not yet reaching full cell death and detachment. This was an interesting observation since PYO has been reported to reach concentrations as high as 100 μM in the lung.^7^ While circulating concentrations are likely to be much lower and, therefore, below the toxicity threshold, it is plausible that they could reach the 1 to 5 μM at the liver, the main site of xenobiotic metabolism.

**Figure 4.**
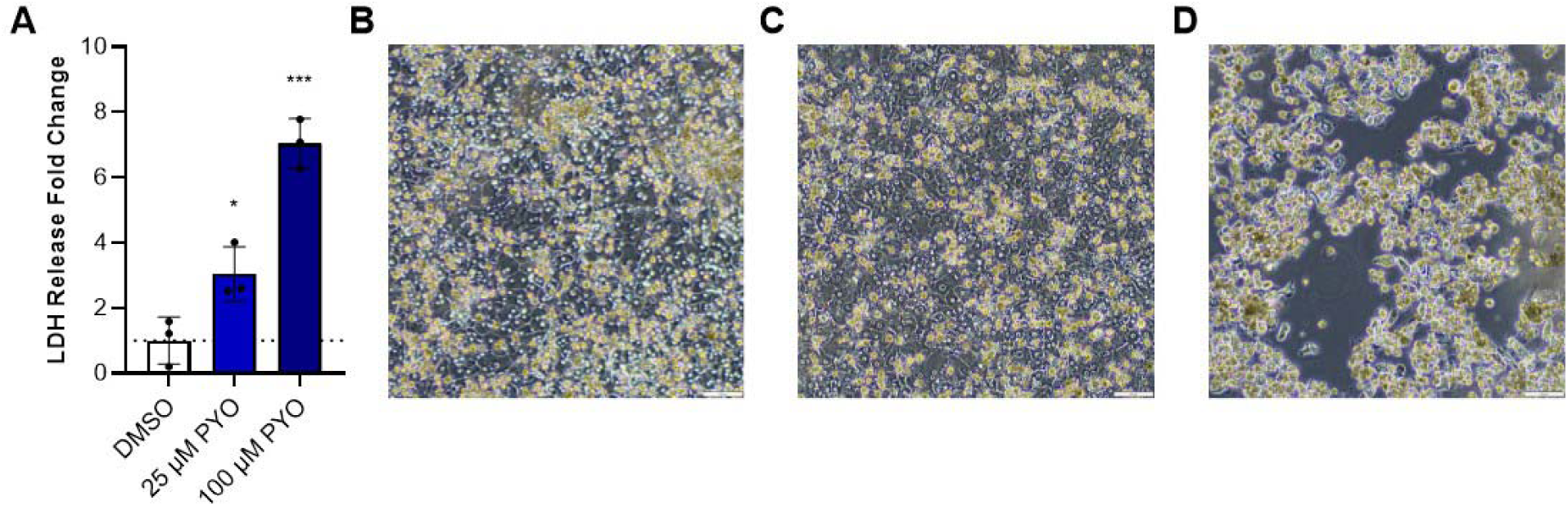
Hepatocyte cytotoxicity of PYO at 25 and 100 μM. *A*, Normalized LDH release at 72 h for primary hepatocytes treated with 0.1% DMSO, 25 μM and 100 μM PYO. Bright field microscopy images of hepatocytes at 72 h treated with 0.1% DMSO control (*B*), 25 μM (*C*) and 100 μM PYO (*D*). Scale bar is 50 µm. The bar graphs represent the means of three biological replicates (individual data points shown) with error bars as standard deviations.

Beyond the direct cytotoxicity to the host cells, certain bacterial secondary metabolites may have evolved to provide protection to the pathogen by hijacking endogenous host defenses and co-opting them to more expeditiously clear molecules that are directly toxic to the pathogen. As such, this may represent an underappreciated pathogen survival mechanism. Indeed, with their myriads of secondary metabolites, pathogenic microorganisms have developed mechanisms to make the host local environment more hospitable for colonization and proliferation.^3–5, 11^ The phenomenal multi-function of the secondary metabolites secreted by these organisms in their endeavors is only beginning to be unveiled. Further efforts to understand how pathogens utilize secondary metabolites to manipulate host metabolism are currently underway by our lab and others.

## MATERIALS AND METHODS

### Reagents

PYO was purchased from Millipore Sigma as a 5 mg/mL ready-made, 0.2 µm filtered solution in DMSO (purity 99.9% determined by HPLC). MB and RIF were also obtained from Millipore Sigma (≥ 96% purity). Plateable, cryopreserved primary human hepatocytes from a ten-donor mix gender pool (Cat #: HPCH10^+^) and corresponding reagents (OptiThaw, OptiPlate, OptiCulture, and OptiMatrix) were purchased from XenoTech, LLC. Ciprofloxacin and oxociprofloxacin were purchased from ThermoFisher and Santa Cruz Biotechnology, respectively, and desythelene ciprofloxacin and ciprofloxacin-d_8_ were acquired from Cayman Chemical (all > 96% purity).

### Induction of *CYP1A2* Expression by PYO in Primary Adult Human Hepatocytes

Hepatocytes were thawed and seeded onto collagen-coated plates at 1.3 × 10^6^ cells/mL according to manufacturer protocols and maintained in a humidified incubator at 37 °C and 5% CO_2_. After 24 h, the plated human hepatocytes were treated once daily for two or three days with medium containing PYO at different concentrations, in biological quadruplicate. An initial test at 25 and 100 µM PYO and 0.1% DMSO was conducted and LDH release was measured at 72 h. Follow up experiments included treatments at 1, 5, and 20 µM PYO, DMSO solvent control (0.1%), and MB (10 µM, known CYP1A2 inducer) and/or RIF (10 µM, known CYP3A4 inducer) as positive controls. qPCR was performed at 48 h, and metabolic activity was measured at 72 h as described below. During the entire time of the assays, the health of the hepatocytes was monitored via visual observation of the monolayer, and the LDH release assay was carried out to measure cytotoxicity (as described below).

At the end of the 72-h treatment period, ciprofloxacin metabolites were monitored by liquid chromatography tandem mass spectrometry (LC-MS). Ciprofloxacin, dissolved in DMSO, was added to the culture plates, in quadruplicate, to yield a final concentration of 30 µM and incubated for up to 120 min. Aliquots of the ciprofloxacin incubations (50 µL) were taken and transferred to an equal volume of methanol (containing 40 ng/mL ciprofloxacin-d_8_ as internal standard). After centrifugation, supernatants were analyzed by LC-MS for formation of ciprofloxacin metabolites.

### qPCR Assays

Hepatocytes were lysed and assayed for mRNA levels using the following qPCR procedure. Briefly, the primary human hepatocytes were processed for mRNA isolation using the Qiagen RNeasy mini kit. cDNA was synthesized with the BioRad iScript Reverse Transcriptase kit and qPCR was performed in duplicate with SYBR green primers (supplemental **Table S1**, Integrated DNS Technologies) and SsoAdvanced supermix (BioRad) on a Bio-Rad CFX96 instrument. Fold change in mRNA level was calculated by the ΔΔCt method using *GAPDH* as reference gene. The mRNA ratio between the treatment and DMSO control groups was calculated to obtain the induction fold change.

### LC-MS Analysis of Ciprofloxacin Metabolite Formation

Supernatants were analyzed by LC-MS for metabolite formation with ciprofloxacin-d_8_ as an internal standard using a Waters Acquity UPLC interfaced by electrospray ionization with a Waters Xevo TQ-S micro mass spectrometer (MS) in positive ionization mode with multiple reaction monitoring (MRM) scans. The MS source conditions were as follows: 0.5 kV for the capillary voltage, 150 °C for the source temperature, 500 °C for the desolvation temperature and 900 L/h for the desolvation gas flow. The following mass transitions (including collision energy, CE, and collision voltage, CV) were used: 332>231 (CE = 36 V, CV = 40 V) for ciprofloxacin, 340>249 (CE = 25 V, CV = 40 V) for ciprofloxacin-d_8_, 346>328 (CE = 18 V, CV = 40 V) for oxociprofloxacin, and 306>268 (CE = 22 V, CV = 32 V) for the desethylene metabolite. The analytes were separated on a Waters BEH Phenyl column (1.7 µm, 2.1 × 100 mm) by flowing water and acetonitrile with 0.1% formic acid at 0.3 mL/min and using the following gradient: 2% organic held for 1 min, increased to 20% over 4 min, then increased to 98% over 2 min and held at 98% for 1 min. The MS peaks were integrated using the QuanLynx software (version 4.1, Waters Corp.) and the analyte/internal standard peak area ratios were used for relative quantification of the ciprofloxacin metabolite. In addition, MS/MS spectrum of the oxociprofloxacin metabolite formed in the human hepatocyte incubations was compared to an authentic standard. Due to low metabolite level in the supernatant samples, four replicates of the MB-induced hepatic incubations were pooled and concentrated to obtain the MS/MS spectrum for the metabolite formed in incubations. The same LC-MS conditions described above were used for the daughter scan, except for the daughter *m/z* set at 346 amu and collision energy as 30 V. Retention time and MS/MS spectrum matched the oxociprofloxacin standard.

### Hepatocyte LDH Cytotoxicity Assays

To assess the propensity for PYO to induce cytotoxicity in the hepatocytes, an LDH release assay was conducted using the LDH-Glo Cytotoxicity Assay (Promega). Primary cryopreserved human hepatocytes were treated at 1-100 μM PYO, 10 μM MB, or DMSO vehicle control as described above. For testing activity of the released LDH, aliquots of culture medium were diluted into LDH storage buffer (200 mM Tris-HCl pH 7.3, 10% glycerol, 1% bovine serum albumin), and an aliquot of 50 μL was transferred to a new 96-well plate, in duplicate. The reaction solution (50 μL) from the kit (containing lactate, NAD^+^, reductase, reductase substrate, and Ultra-Glo rLuciferase) was then added and luminescence was measured after 30 min. Luminescence from the media background was subtracted from all wells, and normalized LDH release was calculated compared to the DMSO control.

### Statistics

All results are reported as the mean ± standard deviation. Statistical analyses were performed via GraphPad Prism (version 9.4.1). Differences between the DMSO control and the treatments were determined by t-tests and significance is shown by *, **, *** and **** corresponding to the following levels: p ≤ 0.05, p ≤ 0.01, p ≤ 0.001, and p ≤ 0.0001, respectively.

## Supporting information

Gracey-et-al-Supplemental-File

## Acknowledgment

This work was generously supported by NIH-NIAID grant R01 AI176245 (to J.N.L.).

## Abbreviations used

ADME: Absorption, Distribution, Metabolism, and Excretion
AhR: aryl hydrocarbon receptor
CE: collision energy
CV: cone voltage
DMSO: dimethyl sulfoxide
HPLC: high-performance liquid chromatography
LC-MS: liquid chromatography-mass spectrometry
LDH: lactate dehydrogenase
MB: methylene blue
MRM: multiple reaction monitoring
MS/MS: tandem mass spectrometry
P450: cytochrome P450
PYO: pyocyanin
qPCR: quantitative polymerase chain reaction
RIF: rifampicin

## Supporting Information

The Supporting Information contains:

◾ qPCR primer sequence details, effect of PYO on *CYP3A4* expression and LDH cytotoxicity at low PYO concentrations.

