## Supplementary material for "*Pseudomonas aeruginosa* co-opts host antibiotic metabolism by pyocyanin induction of cytochrome P450 enzymes": Gracey-et-al-Supplemental-File

**Supporting Information**

**Table S1. SYBR Green primers for qPCR.**

| **Gene** | **Primer sequences** |
| --- | --- |
| *CYP1A2* | CCTCCTTCTTGCCCTTCAC |
|  | CAGCTCTGGGTCATGGTTG |
| *CYP3A4* | GGGAAATATTTTGTCCTACCATAAGG |
|  | ATCATGTCAGGATCTGTGATAGC |
| *GAPDH* | ACATCGCTCAGACACCATG |
|  | TGTAGTTGAGGTCAATGAAGGG |

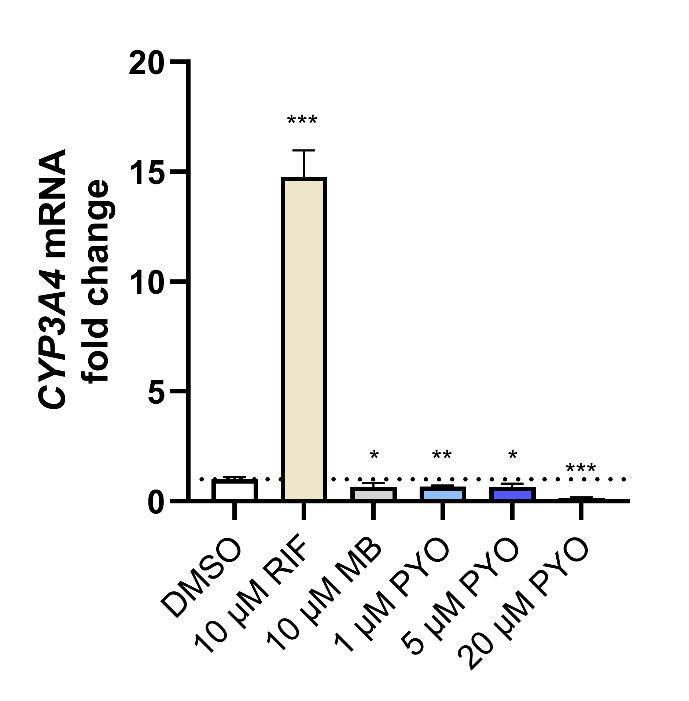

**Figure S1. No PYO effect on *CYP3A4* expression in pooled plated cryopreserved human hepatocytes.** Fold change of *CYP3A4* mRNA in hepatocytes treated once daily for two days with either DMSO (solvent control), 10 µM MB, 10 µM rifampicin (RIF, CYP3A4 positive control for induction) or PYO at 1, 5 or 20 µM. The bar graphs represent the means of four biological replicates (± standard deviation) with individual data points represented. Each biological replicate was assayed by duplicate qPCR reactions and fold induction was calculated via ΔΔCt method (reference gene *GAPDH*).

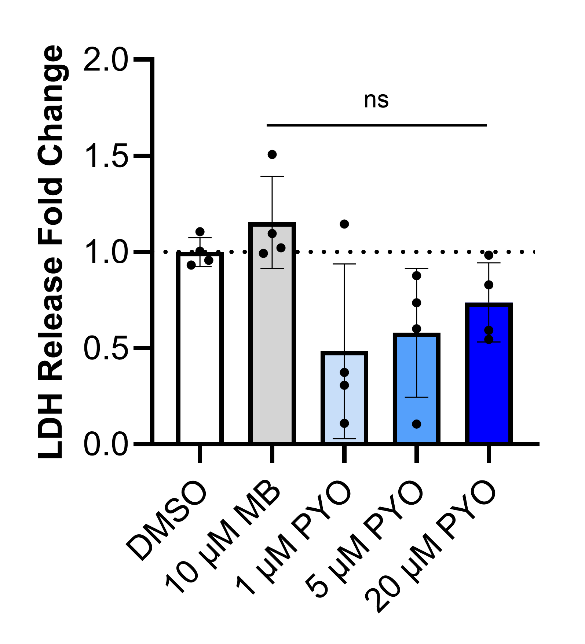

**Figure S2.** **Lactate dehydrogenase (LDH) release results indicated no hepatocyte cytotoxicity of** **PYO at and below 20 μM.** Normalized LDH release at 72 h for primary hepatocytes treated with 0.1% DMSO, 10 μM MB and 1, 5, and 20 μM PYO. The bar graphs represent the means of four biological replicates (± standard deviation) with individual data points represented. ns, not significant.
